# JBrowse Jupyter: A Python interface to JBrowse 2

**DOI:** 10.1101/2022.05.11.491552

**Authors:** Teresa De Jesus Martinez, Elliot A. Hershberg, Emma Guo, Garrett J Stevens, Colin Diesh, Peter Xie, Caroline Bridge, Scott Cain, Robin Haw, Robert M. Buels, Lincoln D. Stein, Ian H. Holmes

## Abstract

**Motivation:** JBrowse Jupyter is a package that aims to close the gap between Python programming and genomic visualization. Web-based genome browsers are routinely used for publishing and inspecting genome annotations. Historically they have been deployed at the end of bioinformatics pipelines, typically decoupled from the analysis itself. However, emerging technologies such as Jupyter notebooks enable a more rapid iterative cycle of development, analysis and visualization.

**Results:** We have developed a package that provides a python interface to JBrowse 2’s suite of embeddable components, including the primary Linear Genome View. The package enables users to quickly set up, launch and customize JBrowse views from Jupyter notebooks. In addition, users can share their data via Google’s Colab notebooks, providing reproducible interactive views.

**Availability:** JBrowse Jupyter is released under the Apache License and is available for download on PyPI. Source code and demos are available on GitHub at https://github.com/GMOD/jbrowse-jupyter.

## 1 Introduction

Genome browsers are popular tools for displaying genome annotations [1]. Web-based browsers, particularly JavaScript tools such as IGV.js [2] and JBrowse [3], allow for a more responsive user experience by performing computation on the client, reducing server and network load. These genome browsers are versatile and widely-adopted [4].

Although JBrowse is straightforward to deploy, the need to administer a web server could present a barrier to entry for some users, particularly those more comfortable working within Python or R. Motivated by this, we recently released JBrowseR, a version of JBrowse designed to run in an R environment or a Shiny app [5]. This was enabled by the React-based architecture of JBrowse 2, which makes possible the development of embeddable components for distinct JBrowse views.

Python, similarly to R, is one of the most widely used programming languages in computational biology. The Jupyter notebook, a web-based interactive environment for Python (and other languages), is one of the leading open source platforms for data analysis [6, 7]. Several tools exist for interacting with genome browsers in Python. D3GB and IGV.js’s integration with Dash provide interactive linear displays of the genome whereas pygbrowse provides static snapshots [2, 8, 9]. However, these tools have limited extensibility or interactivity.

To broaden the options for users of genome browsers in Python environments, we created JBrowse Jupyter, a Python package for configuring, creating, and launching JBrowse views from Jupyter notebooks and other Python applications. JBrowse Jupyter uses the Dash framework, a bridge from React components to Python code [10]. The JBrowse Jupyter package currently allows users to embed two of the most popular JBrowse 2 views: the Linear Genome View and the Circular Genome View. The package aims to make it easier for users to configure genome browsers in Jupyter notebooks and to enable interactive exploration and visualization of genomic data from within Python.

## 2 Methods

JBrowse Jupyter makes use of Dash JBrowse, a collection of Dash components for JBrowse 2 views, to render React components inside Jupyter notebooks. Dash wraps JBrowse’s React embeddable components (such as the Linear Genome View) for use in any Python Dash environment. JBrowse Jupyter’s integration with Dash and Dash callbacks makes it possible to control the state of the JBrowse views from the controlling Python application. Python code can control a JBrowse instance by using the JBrowseConfig API. Examples of supported operations include updating an assembly for viewing, adding tracks, enabling text searching, and changing default view settings. JBrowse Jupyter supports loading track data from files or from Python data frames. Most genomic data file formats are supported.

Users can take advantage of the existing tools available for Jupyter notebooks. Visual Studio Code provides a Jupyter extension that allows users to create and run cells with the embedded genome browser in their development environments. With Binder and Colab, the community is able to share reproducible notebooks with configured JBrowse views [11, 12].

The package is distributed by the Python Package Index and is compatible with Python 3.6 and higher. The project follows continuous integration and continuous delivery practices that enable automated deployments. Flake8 and Pytest support automatic testing and linting to ensure code quality and Sphinx automatically generates documentation.

## 3 Discussion

JBrowse Jupyter is a flexible tool that reduces the effort required to embed an interactive genome browser in a Jupyter notebook: a user can launch a JBrowse View with fewer than ten lines of code (Fig1).

**Figure 1:**
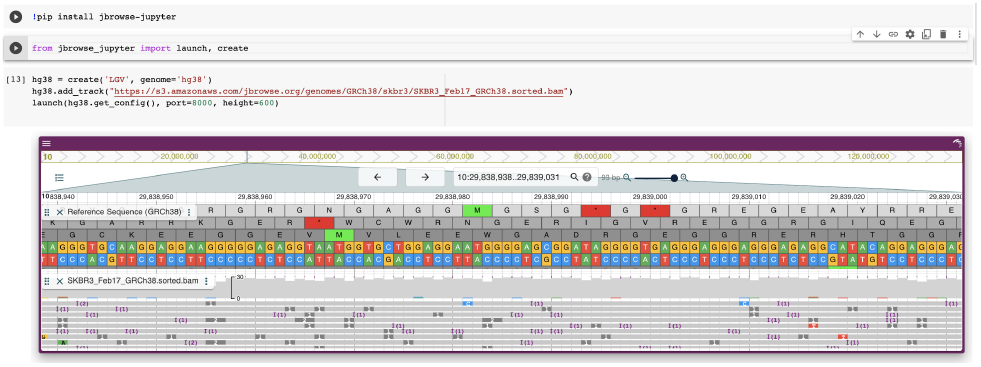
A close-up of the human genome is displayed by JBrowse 2 in a Colab notebook.

We provide several Python Dash applications and Jupyter notebooks along with source code to show the versatility of JBrowse Jupyter. The browser.ipynb notebook demonstrates how to add a track from a Pandas Data frame, add tracks in bulk from distinct data files, and specify initial location in the configuration of the Linear Genome View.

To illustrate application to real data, we provide the skbr3.ipynb Jupyter notebook that visualizes genome sequencing and annotation data from the SK-BR-3 cell line [13]. This demo shows how to create an application that allows users to navigate around genomic locations involved in gene fusions.

## 4 Availability

The package is available for download on PyPI. Source code and demos can be found at https://github.com/GMOD/jbrowse-jupyter.

